# Epigenetic regulation of miR-218 and its host gene, SLIT2 in breast cancer

**DOI:** 10.1101/046482

**Authors:** Ganesh Babu, Arivudai Nambi

## Abstract

Numerous studies have shown that microRNAs are deregulated in various types of cancer. Although miR-218 aberrant expression has been documented in various cancers, the epigenetic regulation of miR-218 is unclear in breast cancer. Our results demonstrate that treatment of breast cancer cells with demethylating agent, 5-azacytidine, resulted in increased expression of miR-218 and its host gene slit-2, suggesting that they are epigenetically regulated in breast cancer. The ectopic expression of miR-218 sensitize breast cancer cell to doxorubicin treatment. The results suggest that miR-218 is epigenetically regulated along with its host gene and function as a tumor suppressor in breast cancer. miR-218 and SLIT2 may have potential therapeutic value in combination with demethylating drug in breast cancer patients.

## Introduction

MicroRNAs (miRNAs) are evolutionarily conserved post-transcriptional regulators and most of them are conserved from the unicellular algae *Chlamydomonas reinhardtii* to mitochondria, give a clue that they are a vital regulatory element in organisms (Molnár et al., 2007). miRNAs are short non-coding RNA, ~22 nucleotide in size and do not code for any protein. In most of the cell types, they act as a negative regulator of gene expression when they bind to their target genes through watson– crick base pairing between the miRNA ‘seed region’ and sequences commonly located in the 3’ UTR, CDS and 5’ UTR of multiple target mRNAs. When they bind with their target genes lead to repression of that protein-coding message, either by transcript destabilization, translational repression, or both (Lee, 2001).

More than 2588 mature miRNA have been documented in human, which may target about 60% of human genes (mirBase). A single miRNAs can target numerous mRNAs whereas individual mRNAs can be targeted by multiple miRNAs, providing vast regulatory potential (Peter, 2010).

A large body of evidence suggests that widespread disruption of miRNA expression is frequently associated with human cancer. Deregulation of miRNAs has been proposed to contribute to oncogenesis in cancer (Croce, 2009; Kasinski and Frank, 2011). Various mechanisms are involved in alteration of miRNA expression in cancer that includes genetic alterations and epigenetic changes which can influence the processing and maturation of miRNA. miRNAs gene are frequently located in genomic fragile sites as well as in cancer-associated genomic regions that are known to be amplified in cancers (Croce, 2009; Kasinski and Frank, 2011).

miR-218 role in various cancers has been investigated and identified as a tumor suppressor miR. It has been found that miR-218 are down regulated in numerous cancers such breast cancer, brain tumor, oral squamous cell carcinoma, nasopharyngeal carcinoma and bladder cancer (Uesugi et al., 2011; Alajez et al., 2011; Tatarano et al., 2011). Even though miR 218 de-gulation have been observed in various cancers, the mechanisms by which miR-218 is silenced has not been well established yet. In this study, we have identified how miRNA 218 expressions is epigenetically regulated along with its host gene breast cancer.

## Materials and methods

### Cell Lines and 5-azacytidine treatment

Breast cancer cell line, HCC712, was routinely maintained in RPMI 1640 supplemented with 10% FCS at 37°C, 5% CO_2_. The demethylating agent 5-azacytidine was prepared in DMSO and filter sterilized. Cells (5x10^5^) were plated in a 25cm^2^ flask in RPMI 1640 supplemented with 10% FCS. 24h later, cells were treated with 12.5 5µM of 5-azacytidine. The medium was changed every 24 h after treatment. RNA was prepared 72h after treatment using the RNeasy kit (Qiagen) according the manufacturer’s guidelines.

### Quantitative real-time RT-PCR

The expression levels of miR-218 (Assay ID: 000521 [Applied Biosystems]) were analyzed by TaqMan quantitative real-time PCR as described in the product manual and normalized to the expression of RNU48. QPCR for SLIT2, SLC6A1 and BCL11A was performed as described before (Dunwell et al., 2009).

### Transfection of miR-218

The following mature miRNA was used in the present study: mature miRNA, Pre-miR miRNA Precursor (has-miR-218;). miR-218 (P / N: AM17100 [Applied Biosystems) was incubated with OPTI-MEM and Lipofectamine RNAiMax reagent (Invitrogen) as described in product manual. Realtime PCR confirmed transfection efficiency of mi-218. MTT Assays was performed as described in product manual (Catalog Number: TA5355/ rndsystems).

## Results and Discussion

### Genomic Organization miR-218

miR-218 is an intergenic miR, they are expressed by two different miR precursors, hsa- mir-218-1 and hsa-mir-218-2. These precursors are encoded in separate genomic locations, 4p15.31 and 5q35.1 respectively. miR-218-1 is housed in intron of SLIT2 where as 218-2 is resided in intron of SLIT3 (Figure 1).

**Figure 1.**
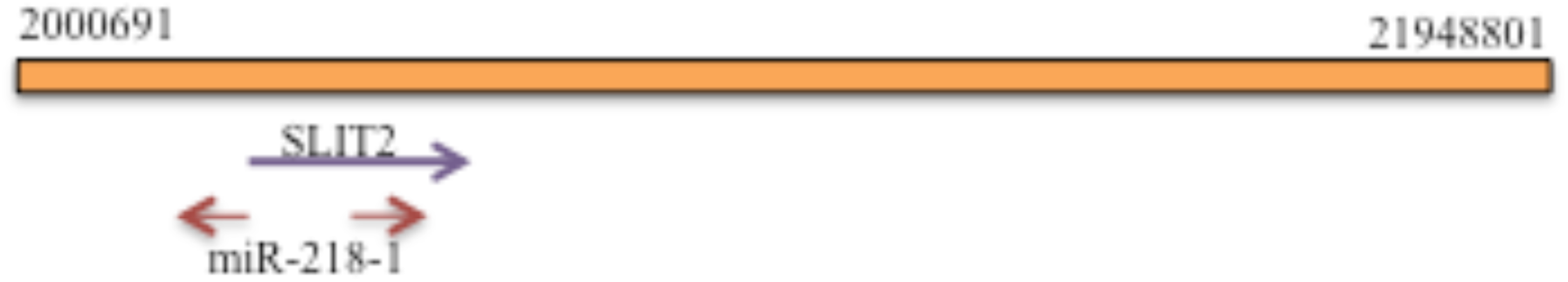
Genomic organization of miR-218-1

As various studies have well documented SLIT 2 locus hyper methylated in several cancers including breast cancer (Alvarez et al., 2010; Kim et al., 2011), we wanted to check that whether the expression miR-218 expression is regulated by hyper methylation. Breast cancer cells were treated with demethylating agent, 5-azacytidine and checked for expression of SLIT2, miR-218 and miR-218 target genes. As shown by Q-PCR, the expression of SLIT2 was significantly induced compared to the control (P<0.001) (Figure 2).

### miR-218 expression and its target genes expression are induced by DNA demethylating agent

QRT-PCR indicated that miR-218 expression in breast cancer cells was strongly elevated than control by 5-azacytidine whereas expression of miR-218 target genes, SLC6A1 and BCL11 A, were reduced significantly ((P<0.001) (Figure 2).

**Figure 2.**
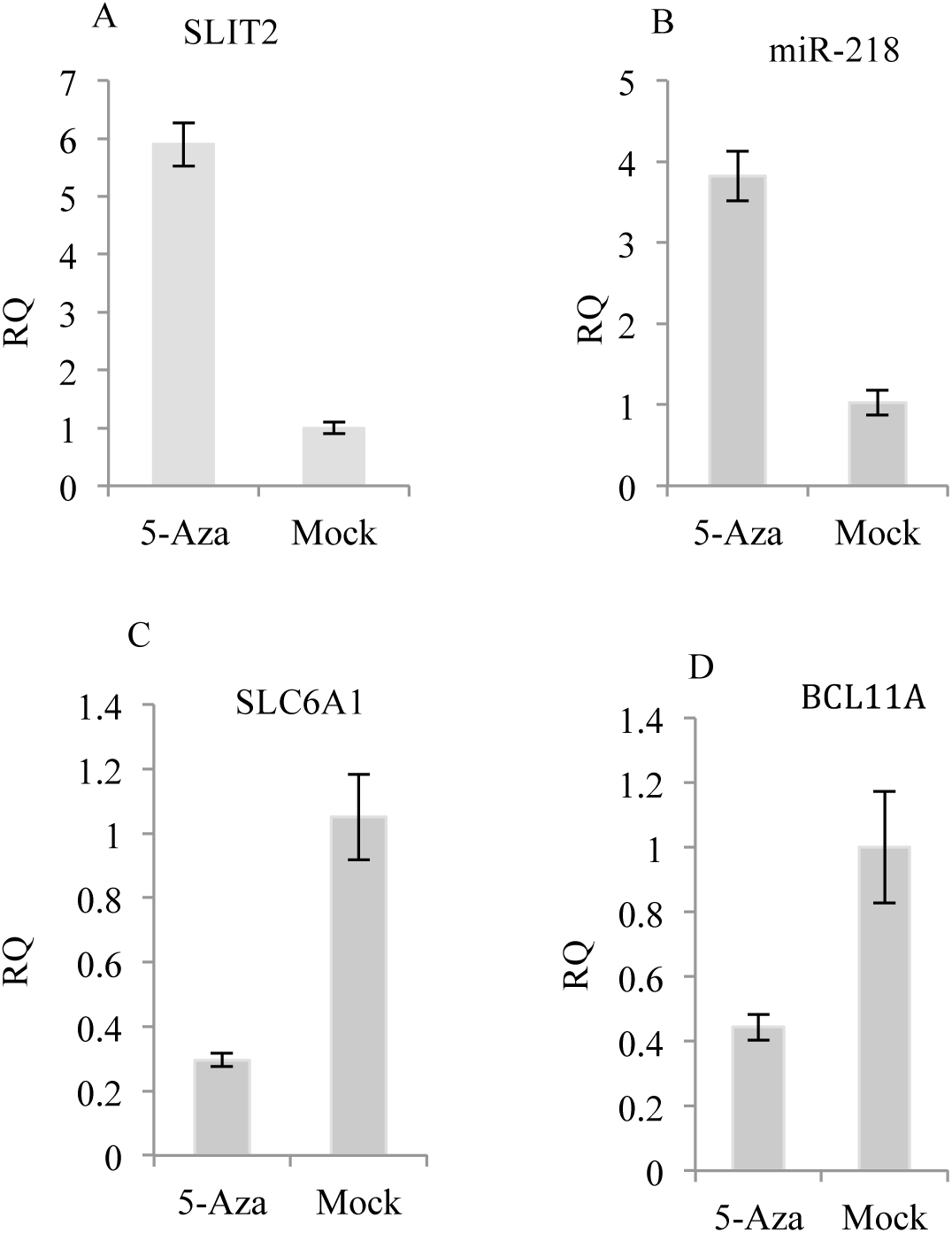
SLIT2 (A) and miR-218 (B) are up regulated in breast cancer cells upon 5-azacytidinetreatment. qRT-PCR analysis for miR-218 target genes, SLC6A1 (C) and BCL11A (D), showing down regulation upon 5-azacytidine treatment.

#### Effect of miR-218 on cell proliferation

To determine the effect of miR-218 on proliferation cancer cells, we overexpressed miR-218 by transfection of miR-218 mimic into breast cancer cell lines. After breast cancer cells were transfected with pre-miR-218, overexpression of miR-218 was confirmed by QRT-PCR. To further investigate, the transfected cells were treated with doxorubicin and cell proliferation was measured after 72h using MTT assays. The results showed that the overexpression of miR-218 significantly reduced cell proliferation in cells transfected with pre-miR-218 cells compared with that in the control cells (Figure 3).

**Figure 3.**
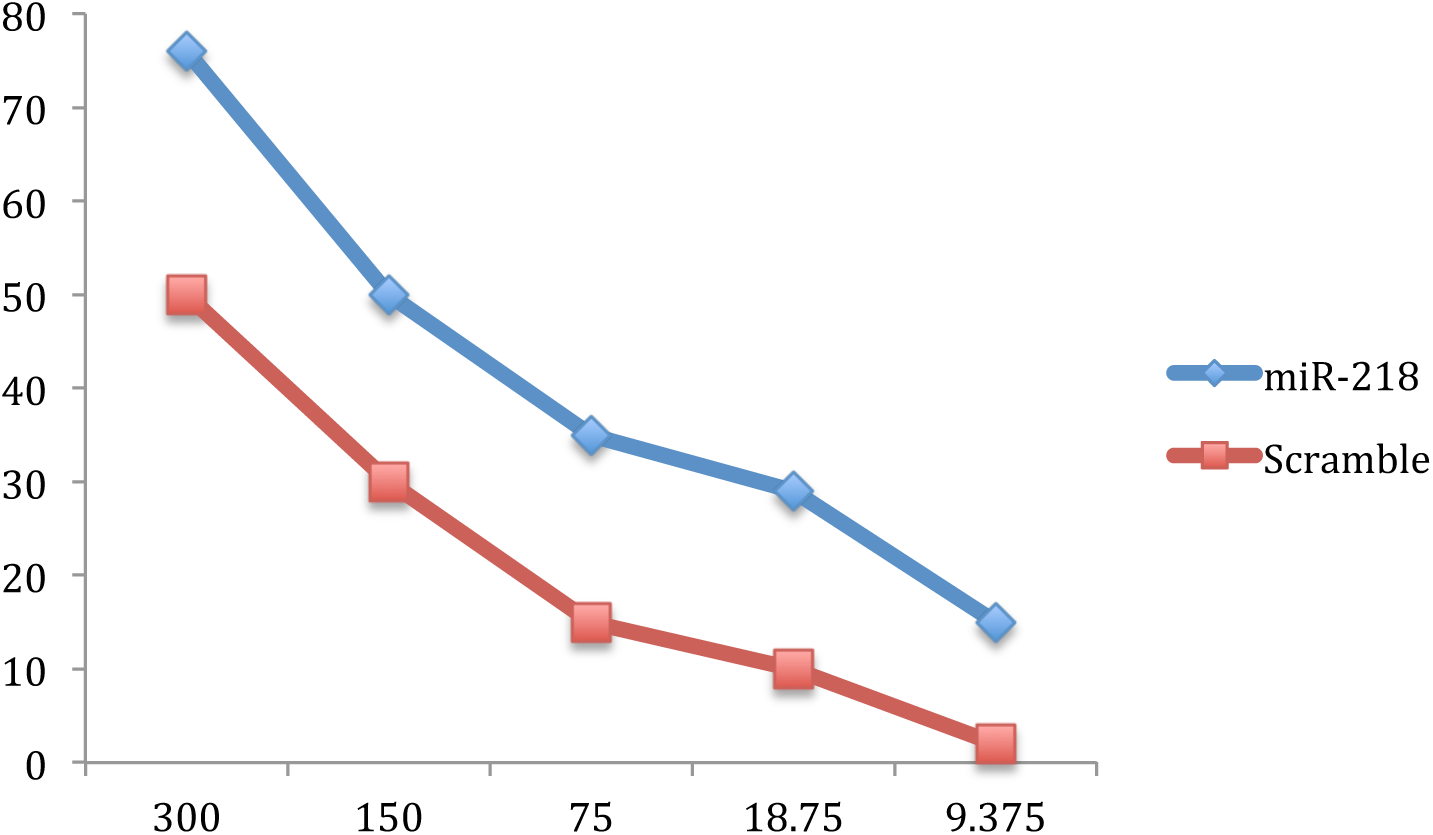
Overexpression of miR-218 potentiates breast cancer cells to doxorubicin treatment, demonstrated as a shift of the miR-218 dose response curves compared to scramble. Cells were plated in 96-well plates at a concentration of 1000 cells per well. After overnight incubation, various concentrations of doxorubicin were added. The MTS assays were performed 72 h after drug treatment. The data were normalized to the doxorubicin-untreated control.

Epigenetic regulation has been shown recently as one of the common inactivation mechanisms of various tumor suppressor genes. Several research teams have shown that promoter methylation of tumor suppressor genes lead to cancer progression. Therefore, in-depth research on the epigenetic regulation on breast cancer is urgently required to improve its early diagnosis and optimize the treatment. SLIT2 gene promoter region has been identified one of the frequently hyper methylated region in lung, pancreatic and breast cancer. In this study we report that miR-218 expression is regulated along with its host gene SLIT2 and SLIT3 through hyper methylation. It has been reported SLIT2 locus is hypermethylated in 36-53% of primary breast tumor. When we treated the breast cancer cells with demethylatiing agent, 5-azacytidine, it strongly induced the expression miR-218 along with its host genes, SLIT2. Meanwhile, 5-azacytidine suppressed the expression of miR-218 target genes SLC6A1 and BCL11A. This observation clearly indicates that expression of miR-218 is controlled by promoter hyper methylation of its hos genes. In addition, ectopic expression of miR-218 in breast cancer cell lines sensitize to doxorubicin treatment. It would be interesting to study further how the ectopic expression miR-218 along with its host genes together affects the cancer progression.

In conclusion, aberrant promoter methylation and associated transcriptional silencing of miR-218 along with its host genes. It suggests that miR-218 along with its host genes, SLIT2 might be amenable to therapeutic intervention by reversal of epigenetic inactivation.

